# Serotonin Engages Divergent 5-HT Receptor Pathways for Cell Type-Resolved Modulation of Prefrontal Layer 5 Microcircuits

**DOI:** 10.64898/2025.12.03.691770

**Authors:** Ramya Rama, Gabriele Radnikow, Danqing Yang, Dirk Feldmeyer

## Abstract

**Background and Purpose:** Serotonin (5-HT) is a key neuromodulator in the prefrontal cortex (PFC), yet its cell type-specific effects across excitatory and inhibitory microcircuits remain incompletely understood. We aimed to determine how 5-HT shapes intrinsic excitability and synaptic transmission in defined neuronal populations of the medial PFC (mPFC).

**Experimental Approach:** We performed whole-cell recordings from morphologically and electrophysiologically characterised layer 5 (L5) neurons in rat mPFC. Pyramidal neurons were classified into adaptive-spiking high input resistance (AS_H_), adaptive-spiking low input resistance (AS_L_), and regular-spiking (RS) types; interneurons were categorised as non-fast-spiking (nFS), regular-fast-spiking (rFS), and stuttering-fast-spiking (sFS) interneurons. These cell types exhibited distinct membrane properties, firing patterns, and axonal projections. We assessed serotonergic modulation using bath-applied 5-HT and subtype-specific receptor mechanisms.

**Key Results:** 5-HT produced divergent postsynaptic responses across pyramidal neuron subtypes: ASH and RS neurons depolarised via 5-HT_2A_ receptor (5-HT_2A_R) activation, whereas AS_L_ neurons exhibited 5-HT_1A_R-mediated hyperpolarisation. Among interneurons, rFS and sFS cells depolarised through ionotropic 5-HT_3A_Rs, while nFS interneurons were largely unaffected. At the synaptic level, 5-HT suppressed excitatory synaptic transmission between pyramidal neurons via presynaptic 5-HT_1B_Rs, and conversely enhanced GABAergic transmission from FS interneurons via presynaptic 5-HT_3A_Rs.

**Conclusions & Implications:** Serotonin exerts bidirectional, cell type-specific serotonergic modulation of local microcircuits that acts to suppress excitation while facilitating inhibition. This balanced and targeted regulation, arising from distinct distributions of somatodendritic and presynaptic 5-HT receptors, provides a mechanistic basis for serotonergic influence on attention, cognitive flexibility, and emotional regulation.

## Introduction

The medial prefrontal cortex (mPFC) plays a pivotal role in higher cognitive functions and emotional regulation, with its activity being profoundly modulated by serotonin (5-hydroxytryptamine, 5-HT) signalling (Goto, Yang et al. 2010, Sakurai and Gamo 2019, Friedman and Robbins 2022). As the primary source of cortical serotonergic innervation, the dorsal raphe nucleus densely projects to the mPFC, where 5-HT receptors are particularly abundant in layer 5 (L5) (Puig and Gulledge 2011). This extensive serotonergic modulation is crucial for maintaining cognitive control and executive function, with its dysregulation implicated in various neuropsychiatric disorders including depression and schizophrenia (Vollenweider and Preller 2020).

In the mPFC, 5-HT modulates neural activity through multiple receptor subtypes that are differentially distributed across neuronal populations (Celada, Puig et al. 2013, Santana and Artigas 2017). While most 5-HT receptors are G protein-coupled, the 5-HT_3A_ receptor forms a ligand-gated cation channel that is permeable to Na^+^, K^+^, and Ca^2+^ ions (Millan, Marin et al. 2008, Nichols and Nichols 2008, McCorvy and Roth 2015). These receptors demonstrate cell-type specific expression patterns: 5-HT_1A_Rs and 5-HT_2A_Rs predominantly localise to dendrites and somata of PNs, while 5-HT_2A_Rs are expressed in the somatodendritic region of specific interneuron (IN) subtypes in the neocortex (Riad, Garcia et al. 2000, Vucurovic, Gallopin et al. 2010). In contrast, 5-HT_1B_Rs are primarily localised to presynaptic axon terminals of both PNs and INs, whereas 5-HT_3A_Rs display selective expression in discrete subpopulations of neocortical INs (Naka and Adesnik 2016, Posluszny 2019).

L5 of the mPFC contains morphologically and electrophysiologically diverse PNs and INs that form specialised microcircuits (van Aerde and Feldmeyer 2015). PNs can be classified into adaptive-spiking (AS) and regular-spiking (RS) subtypes based on their firing patterns, while INs include fast-spiking (FS) and non-fast-spiking (nFS) populations with distinct functional roles (Naka and Adesnik 2016). Although neuronal diversity in L5 has been described in increasing detail, how serotonin (5-HT) regulates the activity of these subtypes remains insufficiently understood, particularly with regard to its effects on synaptic transmission. Clarifying whether 5-HT engages distinct receptor mechanisms to modulate membrane properties and/or synaptic inputs across neuronal subtypes therefore represents an important open question.

To address these questions, we combined electrophysiological recordings with morphological reconstruction to investigate serotonergic modulation of L5 neuronal activity in the mPFC. We determined cell type-specific 5-HT effects on membrane potential in PN and IN subtypes through different receptor pathways. Additionally, we identified a key role for small conductance calcium-activated potassium (SK) channels in mediating 5-HT-induced changes in membrane potential. At the synaptic level, we found that 5-HT_1B_ and 5-HT_3A_ receptors exert opposing presynaptic effects on excitatory and inhibitory transmission. Together, these findings reveal a previously under-appreciated complexity in serotonergic modulation of cortical microcircuits, highlighting cell type-specific regulation of both intrinsic excitability and synaptic efficacy.

## Materials and Methods

### Slice Preparation

The animal experiments were conducted in compliance with FELASA (Federation of European Laboratory Animal Science Association) guidelines, EU Directive 2010/63/EU, and German animal welfare laws. Wistar rats (Charles River, of both sexes) aged 17-21 postnatal days were used, which were anaesthetised with isoflurane, decapitated, and their brains were quickly placed into an extracellular solution containing high Mg^2+^ and low Ca^2+^ concentrations (4 mM MgCl_2_ and 1 mM CaCl_2_), which was chilled and bubbled with 95% O_2_ and 5% CO_2_ to maintain proper oxygenation. The extracellular solution helped to reduce synaptic activity in the brains. The frontal part of the rat brains was attached to a cooled metal stage of a vibration microtome (Leica Biosystems) and cut with a blade at the position indicated in Fig. 2.1. The cut brain tissue was then immersed in the ice-cold extracellular solution and 4-5 coronal slices were cut, each with a thickness of 350 µm. These slices were incubated in the same solution for at least an hour at room temperature (21-24 °C) to allow for recovery. Each individual slice was then transferred to the recording chamber of the patch-clamp setup and perfused continuously (∼5 ml/min) with artificial cerebrospinal fluid (ACSF) containing (in mM): 125 NaCl, 2.5 KCl, 25 glucose, 2 CaCl_2_, 1 MgCl_2_, 1.25 NaH_2_PO_4_ and 25 NaHCO_3_. The perfusion solution was bubbled with 95% O_2_ and 5% CO_2_, and the temperature of the solution in the recording chamber was maintained at a range of 30.5-31.5 °C using a bath heater. Patch pipettes with a resistance ranging from 6 to 10 MΩ were employed to record the data. The patch pipette was filled with an intracellular solution containing (in mM): 135 K-Gluconate, 4 KCl, 10 HEPES, 10 Phosphocreatine, 4 Mg-ATP, and 0.3 GTP (PH 7.4, ∼300 mOsm). Biocytin (5 mg/ml, Sigma, Munich, Germany) was added to the intracellular solution to stain neurons during electrophysiological recording. A different solution containing (in mM) 105 Na-gluconate, 30 NaCl, 10 HEPES, 10 Phosphocreatine, 4 Mg-ATP, and 0.3 GTP was used to search for pipettes during paired recordings.

### Electrophysiological recordings

Bright-field illumination was used to visualise layer borders. L2 was readily identifiable as a thin dark band. L5 was distinguished from L3 and L6 by the large size of pyramidal neuron somata with thick, long apical dendrites. The area of the mPFC was observed to increase from the first slice to the fifth. All neurons were recorded in L5 of the prelimbic cortex. Infrared differential interference contrast (IR-DIC) video microscopy was used for visualisation of single neurons. Pyramidal neurons (PNs) and interneurons (INs) were differentiated by the appearance of their cell bodies and action potential (AP) firing patterns, and post hoc identification was aided by 3D reconstructions of axon and dendritic projections. Whole-cell patch-clamp recordings were performed using an EPC10 amplifier (HEKA, Lambrecht, Germany). The sampling rate for the signals was 10 kHz and electrophysiological signal filtered at 2.9 kHz. Analysis was conducted off-line using the Igor Pro software (Wavemetrics, USA).

To assess the single AP characteristics and active properties of AP firing, a series of 1 s-long current steps, ranging from −100 pA to +500 pA in 10–25 pA stepped depolarisations was injected at 1 s intervals. Continuous recordings of membrane potential changes were made in current-clamp mode with TTX applied to suppress AP generation. Due to the low synaptic connectivity ratio in L5, a modified paired-recording technique incorporating a searching protocol (Method 2; Feldmeyer and Radnikow 2016) was employed. A monosynaptic connection can be found by patching multiple cells in ‘loose cell-attached’ mode. When an AP in the putative presynaptic neuron resulted in a unitary excitatory postsynaptic potential (uEPSP) in the postsynaptic L5 neuron, this presynaptic neuron was repatched with a new pipette filled with biocytin containing internal solution. APs were elicited by current injection in the presynaptic neurons and the postsynaptic response were recorded in whole cell (current clamp) mode, the effects of serotonin on unitary EPSPs were then tested.

### Drug application

All drugs used in this study were obtained from Sigma-Aldrich (Steinheim, Germany) or Tocris (Bristol, UK). Single-cell recordings were conducted in the presence of tetrodotoxin (TTX, 0.5µM). Serotonin (5-HT, 10 µM) was applied via bath perfusion for 200–300 s or puff-applied (100 µM) for 1 second during whole-cell patch-clamp recordings. Baseline measurements were taken for 200–300 seconds, followed by bath application of 5-HT, CGS-12066B (CGS, 5µM) or meta-Chlorophenylbiguanide (mCPBG, 30µM) for 200–300 seconds. A washout or antagonist phase using ketanserine (KET, 3µM) or SB-216641 (SB, 5 µM) lasting 300–600 seconds was then recorded. Paired recordings were performed in the absence of TTX. During paired recordings, 5-HT and/or a receptor agonist/antagonist were bath-applied for 200–400 seconds.

### Histological staining

Following single-cell or paired recordings, brain slices containing biocytin-filled neurons were fixed for a minimum of 24 hours at 4 °C in 100 mM phosphate buffer solution (PBS, pH 7.4) containing 4% paraformaldehyde (PFA). The slices were then rinsed several times with 100 mM PBS and treated with 1% hydrogen peroxide (H_2_O_2_) in PBS for approximately 20 minutes to minimise endogenous peroxidase activity. Next, slices were repeatedly rinsed with PBS and incubated for one hour at room temperature in 1% avidin-biotinylated horseradish peroxidase, containing 0.1% Triton X-100 (Vector ABC staining kit, Vector Lab. Inc., Burlingame, USA). To visualise the biocytin-labelled neurons, the enzymatic reaction was initiated by adding 0.5 mg/ml of 3,3-diaminobenzidine (DAB; Sigma-Aldrich, St. Louis, MO, USA) as a chromogen. This reaction, catalyzed by the horseradish peroxidase (HRP), resulted in the formation of an insoluble brown precipitate that labelled the neurons. The slices were then washed with 100 mM PBS and gradually dehydrated using ethanol in increasing concentrations, followed by immersion in xylene for 2-4 hours. Finally, the slices were embedded using Eukitt medium (Otto Kindler GmbH, Freiburg, Germany).

### Morphological 3D Reconstructions

Computer-assisted 3D morphological reconstructions of biocytin-filled neurons were created using NEUROLUCIDA® software (MicroBrightField, Williston, VT, USA) in combination with an Olympus BV61 microscope at 1000x magnification (100x objective, 10x eyepiece). Neurons were selected for reconstruction based on the quality of biocytin labelling when background staining was minimal. The cell body, dendritic, and axonal branches were manually reconstructed under constant visual inspection to detect thin and small collaterals. Layer borders, pial surface, and white matter were delineated during reconstructions at lower magnification. The tissue shrinkage was corrected using correction factors of 1.1 in the x-y direction and 2.1 in the z-direction (Marx, Gunter et al. 2012). Analysis of 3D reconstructed neurons was done with NEUROEXPLORER® software (MicroBrightField Inc., Willston, VT, USA).

### Data analysis

Electrophysiological signals were analysed using custom-written macros for Igor Pro 6 (WaveMetrics, Lake Oswego, USA). Neurons with a series resistance exceeding 50 MΩ or with a depolarised membrane potential (> −50 mV) after rupturing the cell membrane were excluded from data analysis. The resting membrane potential (V_m_) of the neuron was measured directly after breakthrough to establish the whole-cell configuration with no current injection. The input resistance was calculated as the slope of the linear fit to the current–voltage relationship. For the analysis of single spike characteristics such as threshold, amplitude, and half-width, a step size increment of 10 pA for current injection was applied to ensure that the AP was elicited very close to its rheobase current. The spike threshold was defined as the point of start of acceleration of the membrane potential using the second derivative (d^2^V/dt^2^), that is, using 3x standard deviation of d^2^V/dt^2^ as cut-off point. The spike amplitude was calculated as the difference in voltage from AP threshold to the peak during depolarisation. The spike half-width was determined as the time difference between rising phase and decaying phase of the spike at half-maximum amplitude. The AHP amplitude was determined as the difference in voltage from AP threshold to maximum deflection of the repolarisation. Inter spike interval (ISI) was measured as the average time taken between individual spikes at the current step that elicited close to 10 APs. The adaptation ratio was measured as the ratio of the 10^th^ ISI and the 3^rd^ ISI (= ISI_10_/ISI_3_). The frequency-current (f/I) slope was defined as the slope of the linear fit of the frequency-current response curve within the range of 0 to 300 pA.

Synaptic properties were analysed using previously described methods (Feldmeyer, Egger et al. 1999, Feldmeyer and Radnikow 2016, Qi and Feldmeyer 2016). The properties analysed included the EPSP amplitude, rise time, latency, decay time constant, paired-pulse ratio (PPR), coefficient of variation (CV), and failure rate.

Datasets with more than 10 points were visualised using violin plots with overlaid box plots and individual data points, while datasets with 10 points or fewer were shown as bar graphs with individual data points. Wilcoxon Mann-Whitney U test was performed to assess the difference between individual groups. To assess the differences between two paired groups under different pharmacological conditions, Wilcoxon signed-rank test was performed. Statistical significance was set at *P < 0.05, with **P < 0.01, ***P < 0.001 and ****P < 0.0001 indicating higher levels of significance. Results were considered non-significant when P > 0.05 (denoted as ‘ns’ or ‘not significant’). The value of n indicated the number of neurons/synaptic connections analysed.

## Results

### Electro-Morphological Classification of Layer 5 Pyramidal Neurons

An initial classification of L5 pyramidal neurons in the mPFC was performed by analysing their action potential (AP) firing patterns. Based on their adaptation ratio of the AP firing pattern **(Figure 1A)**, L5 pyramidal neurons (PNs) were classified into two different groups: adaptive-spiking (AS) PNs and regular-spiking (RS) PNs (van Aerde and Feldmeyer 2015). In AS PNs, the inter-spike-interval (ISI) during a spike train increases, resulting in an adaptation ratio (ISI_3_/ISI_10_) smaller than 0.8 (0.7 ±0.1, n = 107). In contrast, RS PNs displayed a consistent ISI after the third AP, leading to an adaptation ratio greater than 0.8 (1.0 ±0.1, n = 92) (van Aerde and Feldmeyer 2015) **(Figure 1A)**. AS PNs were further subdivided based on their input resistance (R_in_) into high R_in_ AS (AS_H_) PNs and low R_in_ AS (AS_L_) PNs (238.1 ±76.8 MΩ, n = 45, 42.1% and 154.7 ±35.0 MΩ, n = 62, 57.9%, respectively; van Aerde and Feldmeyer, 2015). Subsequently, the passive membrane properties, single AP properties and firing properties between these three groups were compared **(Figure 1B)**. Compared to AS_H_ and AS_L_ PNs, RS PNs exhibited the lowest R_in_ and the highest rheobase current.

**Figure 1.**
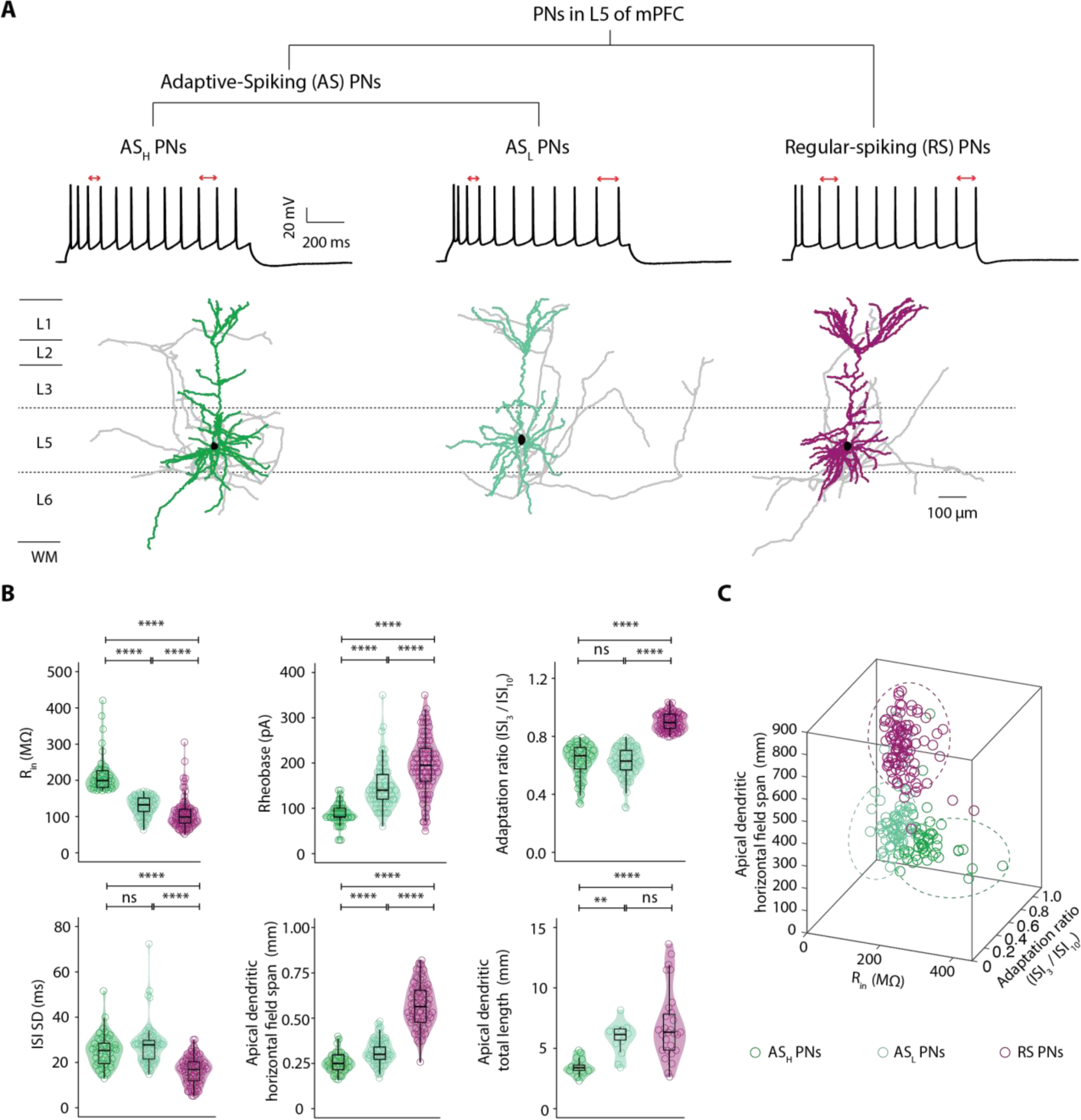
Electrophysiological and morphological classification of L5 PNs in mPFC. **(A)** L5 PNs are classified into adaptive-spiking (AS) and regular-spiking (RS) subtypes based on the adaptation ratio (AdR) of the action potential (AP) firing pattern elicited by depolarising current pulses. AS PNs are further subdivided into high-input resistance (R_in_) AS PNs (AS_H_) and low-R_in_ AS PNs (AS_L_). Representative morphological reconstructions of AS_H_ (left), AS_L_ (middle), and RS (right) PNs are shown, with corresponding firing patterns above each reconstruction. Soma and axon are depicted in black and grey, respectively. Dendrites are colour-coded: dark green for AS_H_, light green for AS_L_, and purple for RS PNs. **(B)** Electrophysiological and morphological properties of AS_H_, AS_L_ PNs, and RS PNs are compared and were presented as violin plots. Colour code as for **(A)**. P < 0.05, **P < 0.01, ***P < 0.001, ****P < 0.0001 for the Wilcoxon Mann–Whitney *U* test; ns, not significant. **(C)** The 3D scatter plot showing the relationship between the adaptation ratio, R_in_, and apical dendritic horizontal field span of AS_H_, AS_L_, and RS PNs. Each circle represents one neuron, and the three PN subtypes are colour-coded as in **(A)**. The three PN groups show a clear separation with little overlap.

Of 199 recorded L5 PNs, 56 neurons were selected for reconstruction based on good staining quality and minimal dendritic truncation, and their morphological properties were analysed. In all AS PNs, the apical dendrite terminated in slender (Sl)-tufted branches (horizontal field span, 0.2 ± 0.1 mm). In contrast, apical dendrites of RS PNs formed broad (Bd), profusely branching apical dendritic tufts (horizontal field span, 0.5 ± 0.1 mm) in L1 **(Figure 1A)**. Within AS PN group, AS_H_ PNs (horizontal field span, 0.2 ±0.1 mm) exhibited more slender tufts compared to AS_L_ PNs, (horizontal field span, 0.3 ± 0.1 mm). In addition, AS_H_ and AS_L_ PNs exhibited relatively strong axonal projections to superficial layers, frequently reaching the pial surface. In contrast, the axons of Bd-tufted RS PNs were observed to project deeply into the white matter. It is important to note that the majority of PN axon collaterals innervate other neocortical regions and subcortical structures and consequently undergo extreme truncation during the slice preparation (Oberlaender, Boudewijns et al. 2011). More electrophysiological and morphological properties of the three types of PNs and their statistical differences are summarised in **Table 1**. To illustrate the distinctions among AS_H_, AS_L_ and RS PNs, a 3D scatter plot was generated, showing the adaptation ratio, R_in_ and horizontal field span of the apical dendrites. Minimal overlap in these parameters was observed between the three groups (**Figure 1C**).

**Table 1:**
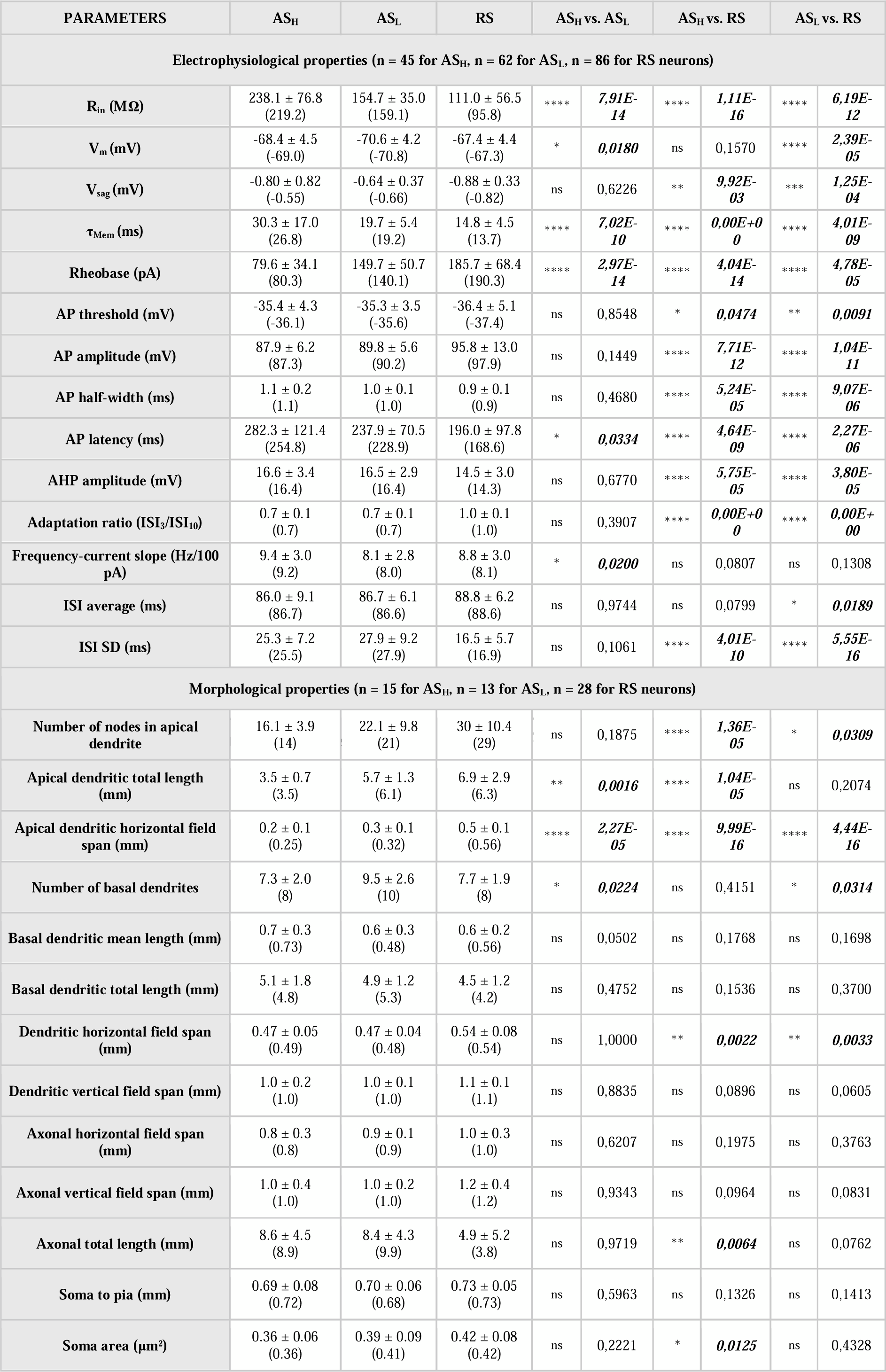
Statistical comparison of the electrophysiological and morphological parameters among L5 AS_H,_ AS_L_ and RS PNs. All properties are given as average ±SD, with median values provided in parentheses. Italic bold font indicates significant differences; *P < 0.05, **P < 0.01, ***P < 0.001, ****P < 0.0001 for Wilcoxon Mann-Whitney U test.

### Electro-Morphological Classification of Layer 5 Interneurons

Interneurons (INs) were grouped into non-fast-spiking (nFS) and fast-spiking (FS) INs based on the adaptation ratio of the AP firing patterns evoked in response to depolarising current pulses (**Figure 2A**). INs with an adaptation ratio of <0.8 were designated as nFS INs (0.6 ±0.2, n = 25), while those with an adaptation ratio >0.8 were classified as FS INs (1.0 ±0.2, n = 54). A subset of FS neurons exhibited irregular firing patterns, characterised by intermittent bursts of action potentials separated by periods of reduced activity, a pattern commonly referred to as stuttering firing (Golomb, Donner et al. 2007, Song, Xu et al. 2013). When compared to regular-FS interneurons (rFS INs), stuttering-FS (sFS) INs showed a larger standard deviation of their inter-spike interval (ISI) (73.0 ±59.3 ms vs 23.0 ± 33.6 ms, P < 0.0001). Moreover, the frequency-current slope (Hz/pA) of rFS INs was larger than that of sFS (80.2 ±52.0 vs. 49.1 ±23.2 Hz/pA, P < 0.05) and nFS INs (80.2 ±52.0 vs. 22.5 ±10.4 Hz/pA, P < 0.0001) (**Figure 2B**). More electrophysiological properties of the three L5 IN types and their statistical differences are summarised in **Table 2**. To differentiate IN subtypes, rheobase, firing frequency and AP half-width were plotted in a 3D scatter plot. The scatter plot illustrates the separation of nFS, rFS and sFS INs, with only little overlap between all types (**Figure 2C**).

**Table 2:**
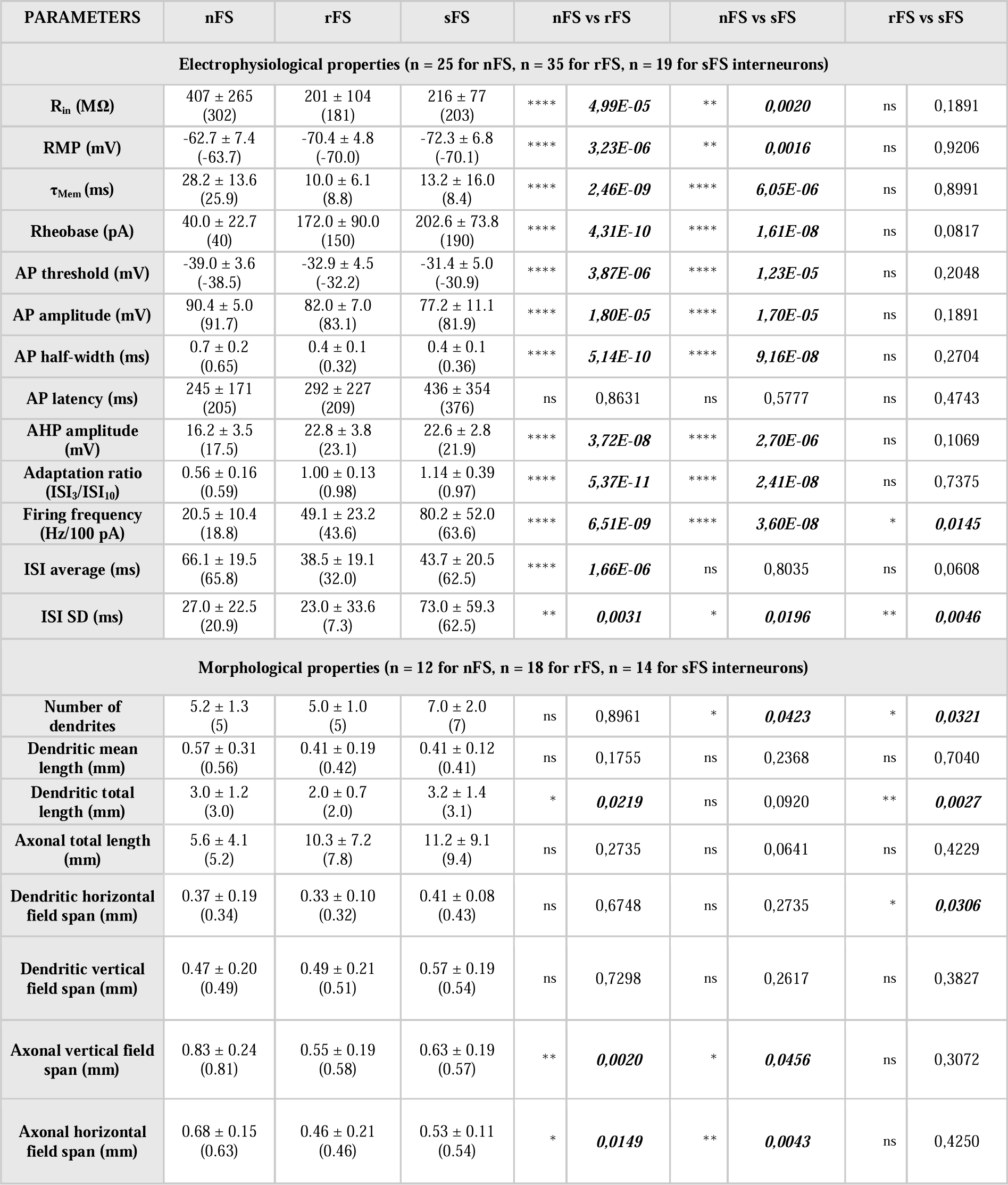
Statistical comparison of electrophysiological and morphological parameters among L5 nFS, rFS and sFS INs. All the axonal and somatodendritic properties are given as average ±SD, with median values provided in parentheses. Italic bold font indicates significant differences; *P < 0.05, **P < 0.01, ***P < 0.001, ****P < 0.0001 for Wilcoxon Mann-Whitney U test.

Of 79 recorded L5 INs recorded, 44 were reconstructed and morphological properties were analysed. INs with large number of axonal truncations and/or an axonal length less than 60% of the average were excluded from the analysis of axonal properties. sFS INs displayed a significantly higher number of primary dendrites compared to rFS (7.0 ± 2.0 vs. 5.0 ± 1.0, P < 0.05) and nFS INs (7.0 ± 2.0 vs. 5.2 ± 1.3, P < 0.05). The axons of nFS INs projected either toward layer 1 or extended into layers 5 and 6, resulting in the largest vertical axonal field span among the L5 IN groups, significantly greater than that of rFS (0.83 ± 0.24 vs. 0.55 ± 0.19 mm, P < 0.01) and sFS INs (0.83 ± 0.24 vs. 0.63 ± 0.19 mm, P < 0.05) (**Figure 2B**). rFS and sFS interneurons exhibited distinct axonal projection patterns: rFS axons extended across layers 5 and 6, whereas sFS axons remained confined to layer 5 (**Figure 2A**). Additional dendritic and axonal parameters for the three IN subtypes, along with their statistical comparisons, are summarised in **Table 2**.

**Figure 2.**
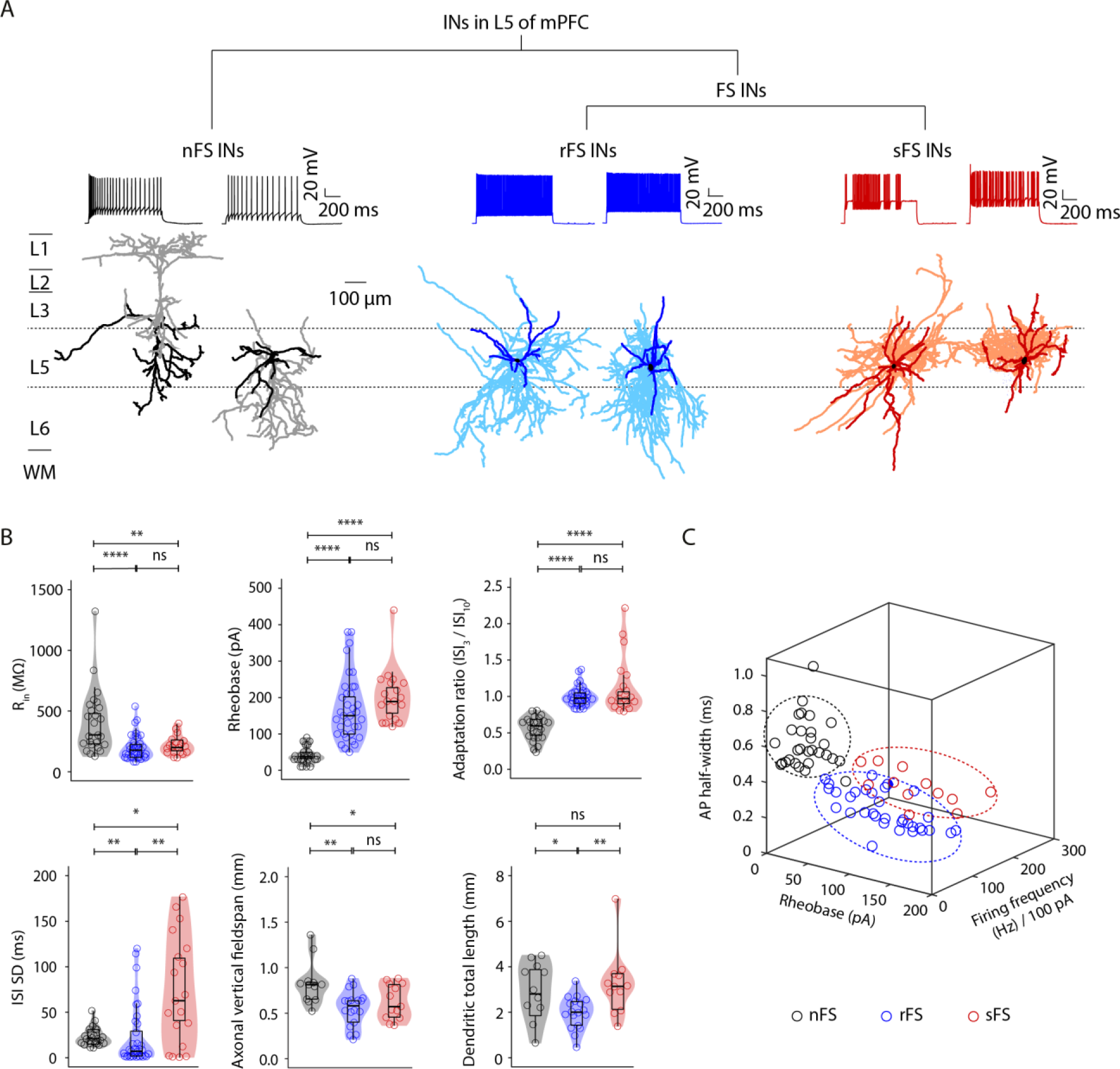
Electrophysiological and morphological classification of layer 5 INs in mPFC. **(A)** Classification of layer 5 (L5) INs based on the AdR of AP trains elicited by depolarising current pulses. L5 INs are grouped into two categories: non-fast-spiking (nFS) and fast-spiking (FS) INs. FS INs are further subdivided into regular-fast-spiking (rFS) and burst-fast-spiking (sFS) INs based on their firing patterns. Representative reconstructions of the three IN types are shown: nFS (left), rFS (middle), and sFS (right), with their corresponding firing patterns illustrated above each reconstruction. Colour coding: nFS INs: somatodendritic domain (black), axon (grey); rFS INs: somatodendritic domain (dark blue), axon (light blue); sFS INs: somatodendritic domain (dark red), axon (light red). **(B)** Summary of the electrophysiological and morphological properties of nFS, rFS, and sFS INs. Colour code as for **(A)**. *P < 0.05, **P < 0.01, ***P < 0.001, ****P < 0.0001 for the Wilcoxon Mann–Whitney *U* test; ns, not significant. **(C)** A 3D scatter plot illustrating the relationship between half-width, rheobase, and firing frequency of nFS, rFS, and sFS INs. Each circle represents an individual neuron, colour-coded as in **(A)**. The three IN subtypes form distinct clusters with minimal overlap.

### Cell type-Specific Modulation of Membrane Potential by Serotonin in Layer 5 Pyramidal Neurons

Next, we investigated how serotonin (5-HT) modulates the membrane potential of L5 PNs. Under current-clamp conditions, 5-HT (10 µM) was bath-applied to L5 PNs in the presence of tetrodotoxin (TTX) to block action potential firing during recordings. In AS_H_ PNs, 5-HT induced a depolarisation of 5.0 ± 2.7 mV. In contrast, AS_L_ PNs showed a hyperpolarisation of – 2.8 ± 1.6 mV in response to 5-HT. In RS PNs, 5-HT also induced a membrane depolarisation, but the effect was significantly smaller than that observed in AS_H_ PNs (2.6 ± 0.7 vs. 5.0 ± 2.7 mV, P < 0.05) (**Figure 3A&B**). In most recorded neurons, 5-HT-induced membrane potential changes were reversible and returned to baseline following washout (**Figure 3A**). To identify the 5-HT receptor subtypes underlying the observed membrane potential changes, specific receptor antagonists were employed. In both AS_H_ and RS PNs, application of ketanserin (KET), a selective 5-HT_2A_ receptor antagonist, during the 5-HT-induced depolarisation completely abolished the response (**Figure 3C**), indicating that 5-HT_2A_ receptors mediate the depolarising effect of 5-HT in these neurons. In AS_L_ PNs, co-application of WAY-100635 (WAY), a selective 5-HT_1A_ receptor antagonist, blocked the 5-HT-induced hyperpolarisation (**Figure 3D**), indicating that this effect is mediated by 5-HT_1A_ receptors.

**Figure 3.**
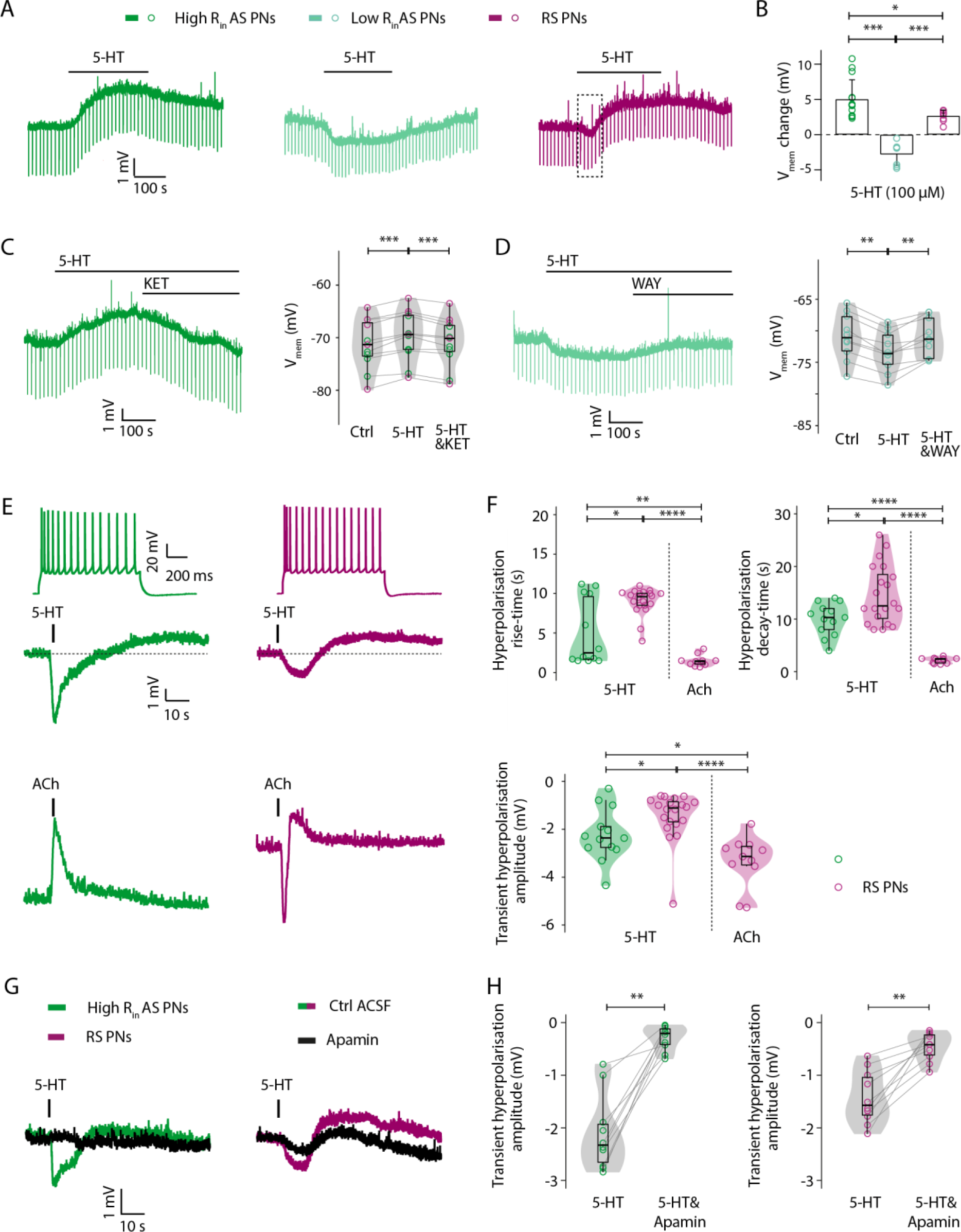
Cell-Type-Specific Modulation of mPFC Layer 5 Pyramidal Neurons by 5-HT: Roles of 5-HT_2A_ and 5-HT _1A_ Receptor Activation. **(A)** Voltage responses of different L5 PN subtypes to bath application of 5-HT (10 µM). AS_H_ PNs (left, green) exhibit membrane depolarisation, whereas AS_L_ PNs (middle, light green) show hyperpolarisation. In RS PNs (right, purple), 5-HT induces a biphasic response with an initial transient hyperpolarisation (dashed box) followed by depolarisation. **(B)** Summary histograms show cell-type-specific 5-HT effects on membrane potential. 5-HT induces greater depolarisation in AS_H_ PNs (n = 14) than RS PNs (n = 9), while AS_L_ PNs (n = 7) exhibit hyperpolarisation. *P < 0.05, ***P < 0.001 for the Wilcoxon Mann–Whitney *U* test. **(C)** Left: application of ketanserine (KET, 3µM), a 5-HT_2A_R antagonist, blocks 5-HT induced depolarisation. Right: violin plots show the effects of 5-HT and KET blockade of 5HT-induced changes (5-HT&KET) on membrane potential in AS_H_ (n = 6, dark green open circles) and RS PNs (n = 5, purple open circles). ***p < 0.001 for the Wilcoxon signed-rank test. **(D)** Left: application of WAY-100635 (WAY, 5µM), a 5-HT_1A_R antagonist, blocks 5-HT induced hyperpolarisation. Right: violin plots show the effects of 5-HT and WAY blockade of 5HT-induced hyperpolarisation (5-HT&WAY) in AS_L_ (n = 10, light green open circles). **p < 0.01 for the Wilcoxon signed-rank test. **(E)** Representative firing patterns (top) of an AS_H_ PNs (dark green) and a RS PNs (purple). Puff application of 5-HT (100 µM for 1s, middle) resulted in a transient hyperpolarisation in both neuron types. In contrast, puff application of ACh (100 µM for 1s, bottom) induced a monophasically depolarisation in AS_H_ PNs while causing a transient hyperpolarisation in RS PNs. **(F)** Violin plots show the effects of 5-HT and ACh on the rise time, decay time, and amplitude of transient hyperpolarisation in AS_H_ PNs and RS PNs. ACh-induced transient hyperpolarisations in RS PNs (n = 11) were larger in amplitude and exhibited faster kinetics than those induced by 5-HT in both AS_H_ PNs (n = 13) and RS PNs (n = 20). *P < 0.05, **P < 0.01, ****P < 0.0001 for the Wilcoxon Mann–Whitney *U* test. **(G)** Pre-incubation of brain slices with the SK-channel blocker apamin reduced the 5-HT-induced transient hyperpolarisation in both AS_H_ and RS PNs. **(H)** Violin plots summarise the differences in apamin-induced reduction of the transient hyperpolarisation in AS_H_ PNs (dark green, n = 12) and RS PNs (purple, n = 10). **p < 0.01 for the Wilcoxon signed-rank test.

It is noteworthy that, in RS pyramidal neurons (PNs), 5-HT induced a biphasic response consisting of an initial transient hyperpolarisation, followed by a sustained depolarisation **(Figure 3A)**. Previous studies have demonstrated that acetylcholine (ACh) can evoke a comparable transient hyperpolarisation in layer 2/3 and layer 5 PNs, that was mediated by SK-type potassium channels (Gulledge and Stuart 2005, Eggermann and Feldmeyer 2009). To better resolve the rapid kinetics of the initial 5-HT response, 100 µM 5-HT was puff-applied for 1 s near the soma of recorded neurons. All RS PNs exhibited a biphasic response consistent with that observed during bath application. In AS_H_ PNs, bath application of 5-HT produced a seemingly monophasic depolarisation; however, puff application revealed a larger and more rapid transient hyperpolarisation compared to that observed in RS PNs **(Figure 3E,F)**. In a separate set of experiments, ACh was also puff-applied to layer 5 PNs to allow direct comparison with 5-HT responses, enabling analysis of the temporal dynamics of the two neuromodulators **(Figure 3E,F)**. Puff application of ACh (100 µM for 1s) induced a monophasically depolarisation in AS_H_ PNs while causing a transient hyperpolarisation in RS PNs (**Figure 3E**). Notably, the ACh-induced transient hyperpolarisation was significantly faster and of greater amplitude than that induced by 5-HT (**Figure 3F**).

We next investigated whether the transient hyperpolarisation induced by 5-HT involved SK channel activation, as previously described for ACh (Gulledge and Stuart 2005, Eggermann and Feldmeyer 2009). To this end, 5-HT was first applied alone to determine amplitude and time course of the transient hyperpolarisation. Subsequently, slices were incubated for 5 minutes in ACSF containing the SK channel antagonist apamin (1 µM). Upon reapplication of 5-HT, the transient hyperpolarisation in AS_H_ PNs was almost completely abolished by apamin (from −2.1 ±0.7 mV to −0.3 ±0.2 mV, P < 0.01). Similarly, in RS PNs, the 5-HT–induced transient hyperpolarisation was significantly reduced in the presence of apamin (from −1.4 ±0.5 mV to −0.5 ±0.3 mV, P < 0.01; **Figure 5G,H**). These findings indicate that SK channels play a key role in mediating the transient hyperpolarising component of the 5-HT response in both AS_H_ and RS neurons.

### Cell Type-Specific Effects of Serotonin on Membrane Potential in Layer 5 Cortical Interneurons

We then examined how serotonin modulates the membrane potential of L5 INs under current clamp conditions. In these experiments, 5 HT (10 µM) was bath-applied in the presence of TTX to prevent action potential firing. In nFS INs, 5 HT did not significantly change the membrane potential (−69.2 ± 8.8 vs. −69.0 ± 9.2 mV, P = 0.1055). In contrast, rFS INs depolarised in response to 5 HT (−68.6 ± 6.0 vs. −61.9 ± 9.9 mV, P < 0.0001). Among sFS INs, most cells (8/12) exhibited a depolarisation, while a smaller subset (4/12) showed a weak hyperpolarisation. Overall, there was no statistically significant change in membrane potential (−75.5 ± 7.4 vs. −72.9 ± 5.8 mV, P = 0.0537) (**Figure 4A&B**).

**Figure 4.**
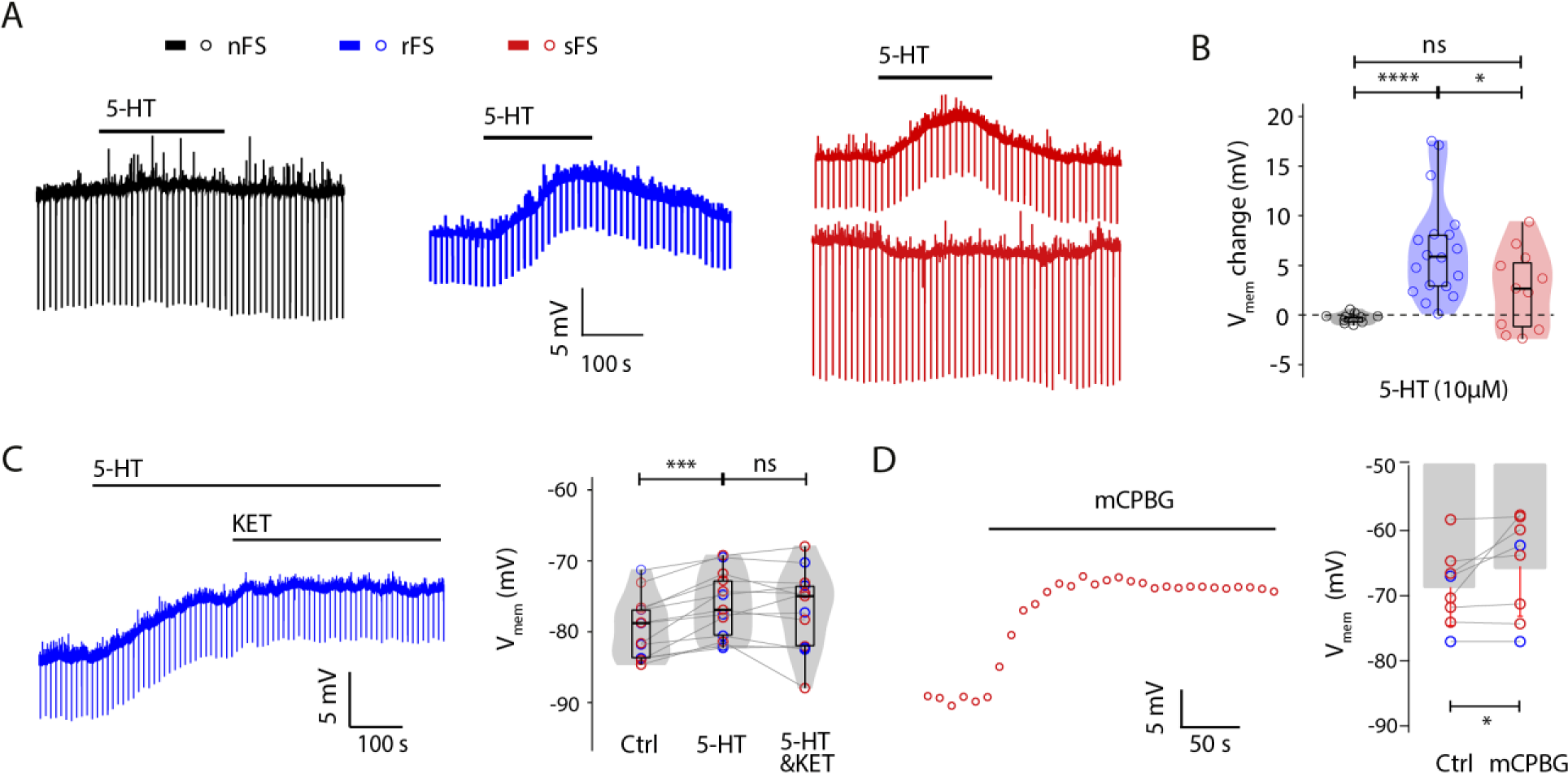
Modulation of Layer 5 Fast Spiking Interneurons in the mPFC by 5-HT through 5-HT_3A_ Receptor Activation. **(A)** Voltage responses of different L5 IN subtypes to bath application of 5-HT (10 µM). 5-HT had no significant effect on the membrane potential of nFS INs (left, black), consistently induced depolarisation in rFS INs (middle, blue), and elicited both depolarising and hyperpolarising responses in sFS INs (right, dark red). **(B)** Violin plots summarise the effects of 5-HT on membrane potential changes in nFS INs (n = 10), rFS INs (n = 18) and sFS INs (n = 11). *P < 0.05, ****P < 0.0001 for the Wilcoxon Mann–Whitney *U* test; ns, not significant. **(C)** Left: the 5-HT induced depolarisation was not affected by administration of KET. Right: violin plots show the effects of 5-HT and co-application of KET and 5-HT on membrane potential in rFS (n = 6, blue open circles) and sFS INs (n = 7, dark red open circles). ***p < 0.001 for the Wilcoxon signed-rank test; ns, not significant. **(D)** Left: bath application of meta-chlorophenylguanidine (mCPBG, 30 µM), a 5-HT_3A_R agonist, induced depolarisation in FS INs. Right: histograms show the effects of mCPBG on membrane potential in FS INs (n = 2 for rFS and n = 6 for sFS INs). *p < 0.05 for the Wilcoxon signed-rank test.

To determine which 5-HT receptor subtypes mediate the observed membrane depolarisations, we applied selective receptor antagonists. In the presence of 5-HT, KET had no consistent effect on the depolarisation of membrane potential (**Figure 4C**). In contrast, application of mCPBG, a 5-HT_3A_ receptor agonist, evoked membrane depolarisations similar to those induced by 5-HT, suggesting that 5-HT_3A_ receptors are primarily responsible for the depolarising effects of 5-HT in L5 interneurons (**Figure 4D**).

### Serotonin Reduces Presynaptic Neurotransmitter Release in Layer 5 Pyramidal Neurons via activating 5-HT1BRs

To elucidate the serotonergic modulation of excitatory synaptic transmission, paired recordings were performed from synaptically coupled mPFC L5 neurons (**Figure 5A&B**). A total of 19 E-E pairs were recorded, comprising ten AS-RS PN pairs, eight RS-RS PN pairs, and one AS-AS PN pair. Since there was no significant differences in uEPSP properties among the different E-E connection types (**Table S1**), the data were pooled and analysed as a single group. Application of 5-HT significantly suppressed synaptic efficacy at L5 E-E connections (**Fig. 5C**). The mean postsynaptic uEPSP amplitude decreased from 1.00 ±0.64 mV to 0.60 ±0.35 mV (n = 15 connections, P < 0.0001), while the paired-pulse ratio (PPR) increased from 0.90 ± 0.22 to 1.19 ± 0.33 (P < 0.0001). In addition, the CV increased from 0.51 ± 0.29 to 0.72 ± 0.33 (P < 0.0001) and the failure rate increased from 13 ± 13% to 27 ± 23% (P < 0.01) at these connections (**Fig. 5C&D**). No significant changes in EPSP latency, 20–80% rise time and decay time were observed, suggesting that 5-HT suppresses synaptic transmission of excitatory connections predominantly via a presynaptic mechanism.

**Figure 5.**
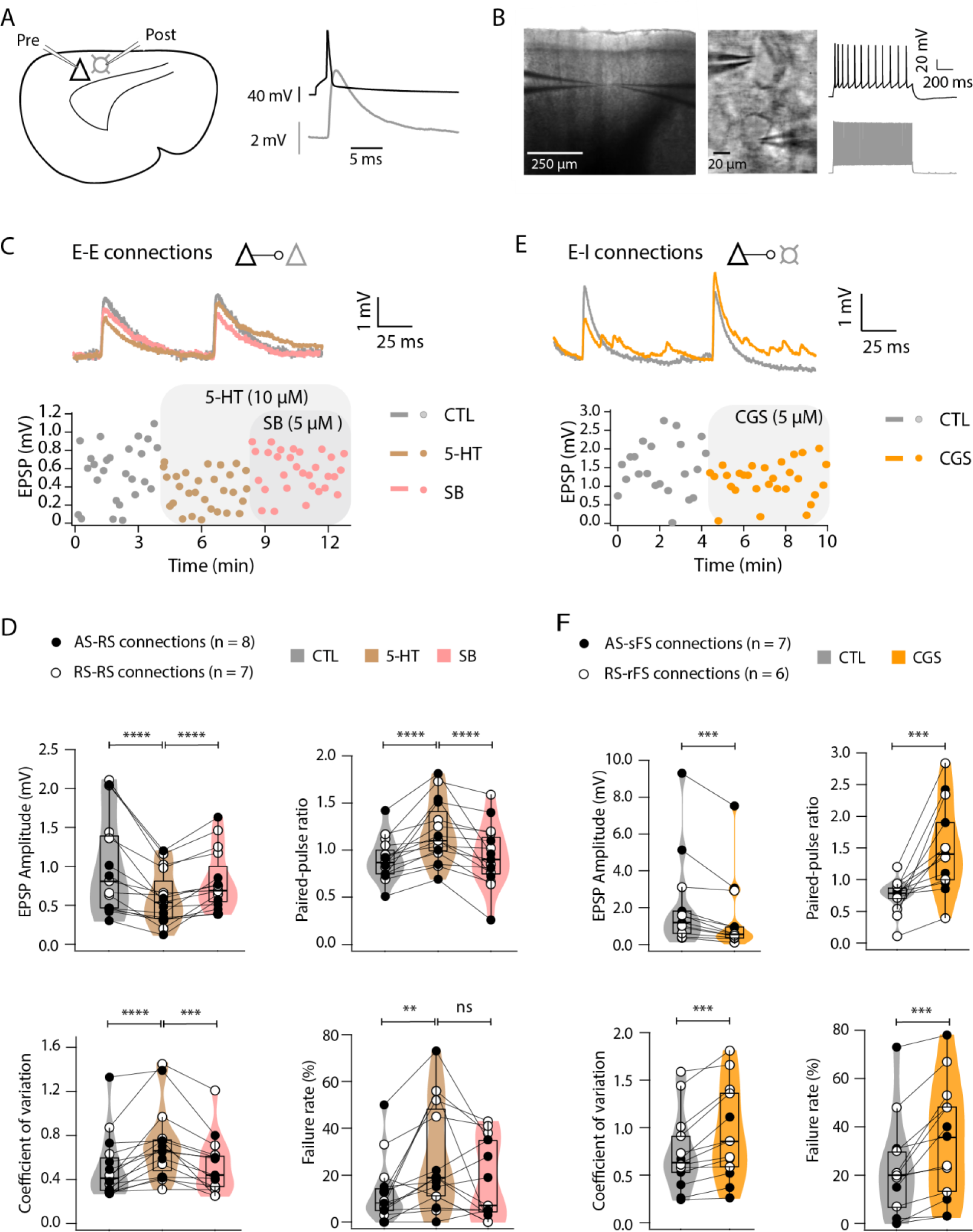
5-HT_1B_ Receptor-Mediated Reduction of Presynaptic Release Probability at Excitatory Synapses. **(A)** Left: schematic representation of a brain slice depicting a pre- and postsynaptic neuron. Right: injection of a current pulse into the presynaptic neuron initiates action potentials (APs, black trace), which in turn evoke an EPSP (grey trace) in the postsynaptic neuron. **(B)** Left: IR-DIC image of a brain slice showing neurons patched in layer 5 of mPFC. Middle and right: higher magnification images (middle) of pre- and post-synaptic neurons along with their firing patterns (right). The presynaptic neuron is RS L5 PN while the postsynaptic neuron is a rFS IN. **(C)** Overlay of the average EPSPs (top) under control condition (grey), during 5-HT application (10 µM, light brown), and of 5-HT and a 5-HT_1B_R antagonist SB-224289 (SB, 5 µM, light pink) from a representative E→E connection in layer 5 of rat mPFC. Bottom, time course of changes in the 1^st^ EPSP amplitude following bath application of 5-HT and of 5-HT and SB (5-HT&SB) in the same E→E pair. **(D)** Violin plots summarise several synaptic properties of E→E connections including EPSP amplitude, PPR, CV and failure rate. L5 AS PN→RS PN connections: black closed circles (n = 8); RS PN→RS PN connections: black open circles (n = 7). **P < 0.01, ***P < 0.001, ****P < 0.0001 for the Wilcoxon signed-rank test; ns, not significant. **(E)** Overlay of the average EPSPs (top) under control conditions (grey) and of 5-HT and a 5-HT_1B_R agonist CGS-12066B (CGS, 5 µM, orange) from a representative E→I connection in layer 5 of rat mPFC. Bottom, time course of changes in the 1^st^ EPSP amplitude following bath application of CGS in the same E→I pair. **(F)** Violin plots summarise several synaptic properties of E→I connections including EPSP amplitude, PPR, CV and failure rate. L5 AS PN→sFS IN connections: black closed circles (n = 6); RS PN→rFS IN connections: black open circles (n = 6). ***P < 0.001 for the Wilcoxon signed-rank test.

**Figure 6.**
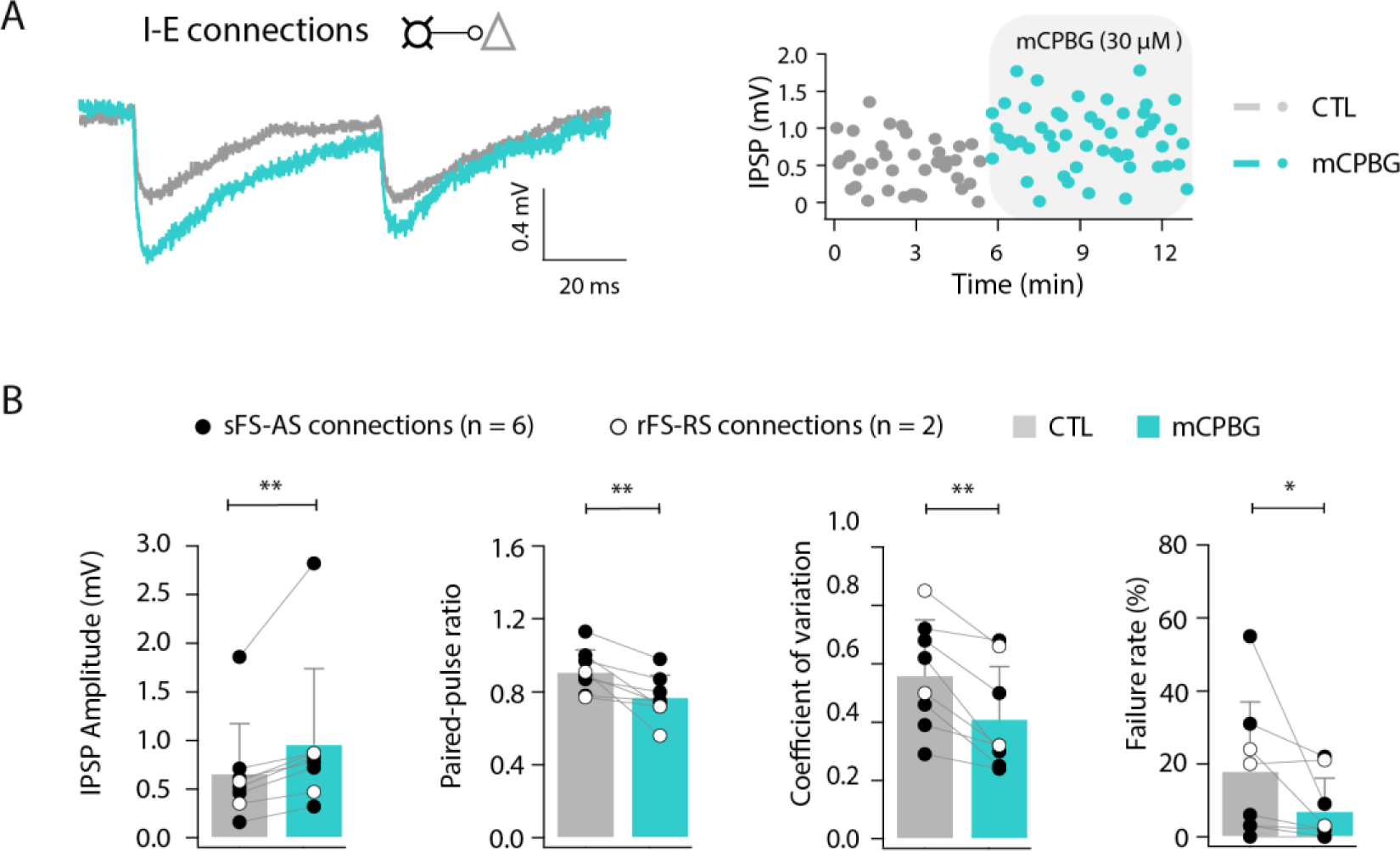
5-HT_3A_ Receptor-Mediated Enhancement of Presynaptic Release Probability at Inhibitory Synapses. **(A)** Left: an overlay of the average IPSPs under control conditions (grey) and during the application of mCPBG (cyan). Right: time course of changes in the 1^st^ IPSP amplitude following bath application of mCPBG in the same I→E pair. **(B)** Summary histograms of several synaptic properties of I→E connections including IPSP amplitude, PPR, CV and failure rate. sFS IN→AS PN connections: black filled circles (n = 6); rFS IN → RS PN connections: black open circles (n = 2). *P < 0.05, **P < 0.01 for the Wilcoxon signed-rank test.

Given that 5-HT_1B_Rs are known to be expressed presynaptically on excitatory terminals in the cortex and can modulate neurotransmitter release (Kjaerby, Athilingam et al. 2016, Madden, Schaefgen et al. 2025), we next tested whether the serotonergic suppression of excitatory synaptic transmission in layer 5 is mediated by 5-HT_1B_R activation. To this end, the selective 5-HT_1B_R agonist SB was co-applied following 5-HT application. The effect of 5-HT on excitatory connections was blocked and uEPSP amplitude recovered from 0.60 ± 0.35 mV to 0.81 ± 0.39 mV (n = 15 connections, P < 0.0001). Furthermore, SB prevented the 5-HT-induced increases in PPR, CV, and synaptic failure rate (**Fig. 5C&D**), indicating that 5-HT reduces neurotransmitter release probability via presynaptic 5-HT_1B_R activation.

We next investigated whether serotonin exerts a similar modulatory effect on excitatory-to-inhibitory (E-I) synaptic transmission. Bath application of the selective 5-HT_1B_R agonist CGS to E-I connections (AS-sFS: n = 6; RS-rFS: n = 7) significantly reduced EPSP amplitude from 2.2 ± 2.6 mV to 1.5 ± 2.1 mV (P < 0.001) and increased PPR from 0.7 ± 0.2 to 1.5 ± 0.7 (P < 0.001). CGS also significantly increased CV and failure rate of E-I connections (**Figure 5E&F**). No significant changes were observed in EPSP rise time, decay time, or latency.

Overall, our results indicate that the suppression of synaptic transmission at both E-E and E-I connections is mediated by a reduced neurotransmitter release probability from presynaptic AS and RS pyramidal neurons in layer 5 of the mPFC. This effect is driven by the activation of presynaptic 5-HT_1_B receptors. A summary of the EPSP properties and statistical comparisons before and after drug application for E-E and E-I connections is provided in **Table S2**.

### Serotonin Facilitates Presynaptic Release in L5 Fast-spiking Interneurons via 5-HT_3A_ Receptors

Next, we investigated how 5-HT influences inhibitory synaptic connections in layer 5 of the mPFC. Neocortical INs express various 5-HT receptor subtypes, including 5-HT_1A_R, 5-HT_2A_R, and 5-HT_3A_R (Celada, Puig et al. 2013, Santana and Artigas 2017). Unlike other 5-HT receptors, 5-HT_3A_Rs are ligand-gated cation channels and are present on approximately 30% of presynaptic IN axon terminals (Koyama, Matsumoto et al. 2000, Ferezou, Cauli et al. 2002). Notably, the presence of 5-HT_3A_Rs has been used in previous studies as a criterion for classifying cortical interneuron subtypes (Rudy, Fishell et al. 2011). To examine the presence and functional role of 5-HT_3A_R activation in inhibitory synapses, we recorded eight inhibitory-to-excitatory (I–E) connections (sFS–AS pairs: n = 6; rFS–RS pairs: n = 2). No inhibitory connections with a presynaptic nFS IN were observed. The selective 5-HT_3A_R agonist mCPBG was bath-applied at a concentration of 30 µM. mCPBG significantly increased IPSP amplitude (0.66 ± 0.51 vs 0.96 ± 0.78 mV, P <0.01), and reduced the PPR (0.91 ± 0.12 vs 0.77 ± 0.12, P < 0.01), CV (0.56 ± 0.19 vs 0.41 ± 0.18, P < 0.01) and failure rate (18 ±19 vs 7 ±9 %, P < 0.05) (**Figure 7A, B**). No significant differences were found in rise time, decay time, or latency. A summary of the IPSP properties and statistical comparisons before and after 5-HT application is provided in **Table S2**.

**Figure 7.**
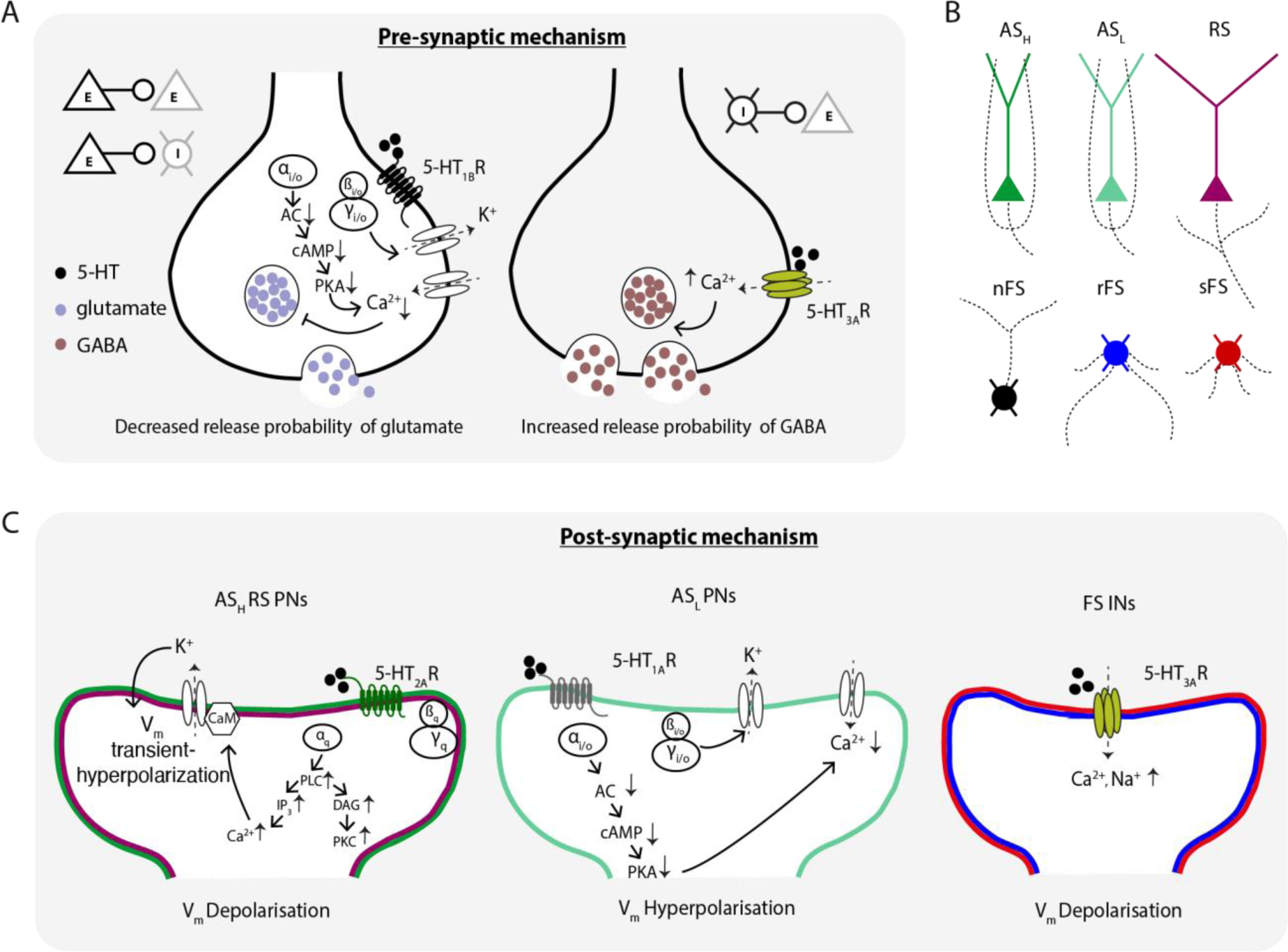
Schematic Representation of Presynaptic and Postsynaptic Modulatory Mechanisms of 5-HT on Different Neuronal Subtypes in Layer 5 of the Rat mPFC. **(A)** Left: 5-HT activates 5-HT_1B_Rs, leading to G_i/o_ protein activation, which inhibits Ca²⁺ influx through downstream signalling, resulting in presynaptic suppression of glutamate release at excitatory synapses. Right: activation of ionotropic 5-HT_3A_Rs by 5-HT directly increases Ca²⁺ influx, enhancing GABA release at inhibitory synapses. **(B)** This schematic panel illustrates the colour coding of excitatory and inhibitory neuronal subtypes in layer 5 of the mPFC. Soma and dendrites are shown in distinct colours, while axons are depicted with black dashed lines, emphasising morphological differences. **(C)** Left: in AS_H_ and RS PNs, activation of 5-HT_2A_Rs induces membrane depolarisation via the G_q_ signalling pathway. The resulting intracellular Ca^2+^ elevation binds to CaM, activating SK channels, which mediate K^+^ efflux, causing an initial brief hyperpolarisation. Middle: In AS_L_ PNs, activation of 5-HT_1A_Rs induces membrane hyperpolarisation via the G_i/o_ signalling pathway. Right: In FS INs, activation of ionotropic 5-HT₃ARs leads to direct Ca^2+^ and Na^+^ influx, resulting in membrane depolarisation. PN and IN subtypes are colour-coded as in **(B)**.

Thus, 5-HT modulates presynaptic neurotransmitter release in L5 neurons of the mPFC via distinct receptor subtypes in a cell type-specific manner. Activation of presynaptic 5-HT_1B_Rs suppresses neurotransmitter release at excitatory synapses, whereas activation of presynaptic 5-HT_3A_Rs enhances release at inhibitory synapses formed by FS INs.

## Discussion

### Diversity of Pyramidal Neurons and Interneurons in Layer 5 of the mPFC

In previous studies L5 pyramidal neurons (PNs) of the medial prefrontal cortex (mPFC) have been classified into two main groups based on their projection targets. The first group consists of L5 PNs that innervate in intratelencephalic (IT) neurons (e.g., in the cortex, striatum, limbic forebrain). L5 neurons of the other group project to extratelencephalic (ET) targets such as the thalamus (Dembrow, Chitwood et al. 2010, Kawaguchi 2017, Baker, Kalmbach et al. 2018, Collins, Anastasiades et al. 2018, Elliott, Tanaka et al. 2018). IT PNs typically exhibit an adaptive-spiking (AS) firing pattern, whereas ET-projecting neurons tend to show regular-spiking (RS) behaviour (Elliott, Tanaka et al. 2018, Moberg and Takahashi 2022). Building on prior classifications, our study further resolves the heterogeneity of AS PNs in mPFC layer 5 by distinguishing two subtypes based on input resistance: high-R_in_ (AS_H_) and low-R_in_ (AS_L_) neurons, following criteria described previously (van Aerde and Feldmeyer 2015). AS_H_ neurons exhibited significantly higher input resistance, greater adaptation ratios, and lower rheobase compared to AS_L_ neurons, indicating higher excitability. Morphologically, AS_H_ neurons possessed more slender apical dendritic tufts with shorter horizontal spread and reduced total apical dendritic length relative to AS_L_ neurons. Despite these differences, both AS_H_ and AS_L_ L5 PN axons projected preferentially to superficial cortical layers, in contrast to RS neurons, which had broader, more extensively branching apical tufts and projected deeply into the white matter.

Here, we identified three distinct interneuron (IN) types in layer 5 of the mPFC based on their action potential firing patterns and axonal morphology. These included non-fast-spiking (nFS) INs and two fast-spiking (FS) subtypes: regular-spiking FS (rFS) and stuttering FS (sFS) INs. The classification was based on electrophysiological properties including spike frequency adaptation, interspike interval variability, AP half-width, and rheobase. Morphological reconstructions revealed that nFS neurons projected to superficial (L1) or deep layers (L5/6), resembling Martinotti cells and deep-layer targeting INs, respectively. In contrast, rFS and sFS neurons exhibited axonal arbors that were consistent with those observed in large and small basket cells, with rFS INs showing a broader vertical axonal span than sFS INs. These findings indicate that a functional and morphological subdivision among L5 FS INs exist that has frequently been disregarded in previous publications (Galarreta and Hestrin 2002, Angulo, Staiger et al. 2003, Goldberg, Jeong et al. 2011, Tremblay, Lee et al. 2016, Chistiakova, Ilin et al. 2019). It also lends support to the hypothesis that even within the PV+ interneuron population, microcircuit-level specialisations exist that can shape the temporal dynamics of inhibition in local cortical networks (Goldberg, Jeong et al. 2011).

### Somatodendritic Effects of Serotonin on Pyramidal Neurons

A previous study reported a dichotomy of PNs in rat mPFC, demonstrating that RS neurons are hyperpolarised, while AS neurons are depolarised by 5-HT (Elliott, Tanaka et al. 2018). In the present study, by further subdividing adapting-spiking neurons into ASH and ASL subtypes based on input resistance, revealing additional diversity within this population. AS_H_ neurons showed a 5-HT_1A_R-mediated hyperpolarisation in response 5-HT application, while AS_L_ neurons exhibited a 5-HT_2A_R-dependent depolarisation. These results suggest that the previously defined AS group may comprise distinct subpopulations differing not only in intrinsic properties but also in their serotonergic receptor complement. Moreover, we found that RS neurons exhibited a biphasic 5-HT response, consisting of an initial transient hyperpolarisation followed by sustained depolarisation. This indicates that serotonergic modulation more complex than previously realised.

The application of specific receptor antagonists was used to identify that the depolarising 5-HT response was mediated by 5-HT_2A_Rs, while the hyperpolarisation in RS and AS_H_ PNs was caused by 5-HT_1A_R activation, consistent with previous studies (Puig and Gulledge 2011, Celada, Puig et al. 2013). A similar receptor-specific effect was reported in rat association cortex, where 5-HT caused either a 5-HT_1A_R-mediated persistent hyperpolarisation or a 5-HT_2A_R-mediated depolarisation leading to AP firing, depending on the PN subtype (Araneda and Andrade 1991). Subtype-dependent responses were also observed in the mouse prelimbic cortex, where 5-HT_2A_R activation induced excitation in commissural/callosal projection neurons, while 5-HT_1A_R activation led to an inhibition of corticopontine projection neurons (Avesar and Gulledge 2012). Co-expression of both receptors within an individual pyramidal neurons is well documented (Martin-Ruiz, Puig et al. 2001, Amargos-Bosch, Bortolozzi et al. 2004, Santana, Bortolozzi et al. 2004, Celada, Puig et al. 2013). However, the net effect of 5-HT varies, likely due to differences in subcellular receptor distribution. 5-HT_2A_Rs are predominantly found in somatic and dendritic compartments (Xu and Pandey 2000, Jansson, Tinner et al. 2001, Mengod, Palacios et al. 2015), whereas 5-HT_1A_Rs are concentrated at the axon initial segment, where they may suppress action potential generation (Azmitia, Gannon et al. 1996, DeFelipe, Arellano et al. 2001, Yin, Rasch et al. 2017). However, somatodendritic expression of 5-HT_1A_Rs has also been reported, suggesting a potential overlap in their spatial distributions (Martin-Ruiz, Puig et al. 2001). These findings underscore the need for further studies to clarify how receptor subtype localisation contributes to the functional diversity of serotonergic responses in cortical PNs.

Both RS and AS_H_ neurons exhibited biphasic responses when 5-HT was delivered via puff application. Interestingly, in RS neurons, the rise and decay of the transient hyperpolarising phase were markedly slower in RS than in AS_H_ PNs. Therefore, during bath application of 5-HT the biphasic response in this neuron type remained undetected due to the low temporal resolution of the applications method. The selective SK channel blocker apamin abolished the hyperpolarisation, indicating the involvement of SK channels (Vogalis, Storm et al. 2003, Kuzmenkov, Peigneur et al. 2022). This response likely results from 5-HT_2A_R–mediated intracellular Ca^2+^ release via the PLC–IP_3_ pathway, which activates Ca^2+^-dependent SK channels (Yamada, Takechi et al. 2004, Hannon and Hoyer 2008, Millan, Marin et al. 2008) (**Figure 7**). The biphasic 5-HT response observed in RS neurons resembles that induced by M1-type muscarinic ACh receptor activation, which also engages SK channels (Gulledge and Stuart 2005, Eggermann and Feldmeyer 2009). Notably, ACh evokes biphasic responses only in RS neurons, and with faster kinetics and larger amplitude—particularly in the mPFC compared to other cortical regions (Gulledge, Park et al. 2007). These differences likely reflect region- and subtype-specific expression patterns of SK channels and G-protein coupled receptors (GPCRs). Together, our findings underscore the importance of neuromodulator timing and delivery mode in shaping neuronal excitability, and point to SK channels and 5-HT_2A_Rs as potential targets for modulating mPFC function in neuropsychiatric conditions.

### Receptor-Dependent 5-HT Responses in Interneurons

L5 PNs of mPFC frequently co-express 5-HT_1A_ and 5-HT_2A_Rs (Santana, Bortolozzi et al. 2004, Puig and Gulledge 2011, Celada, Puig et al. 2013). In contrast, mPFC L5 interneurons exhibit a broader receptor profile that includes the ionotropic 5-HT_3A_ receptor in addition to GPCR-coupled 5-HT receptors, reflecting a greater diversity in 5-HT receptor function in inhibitory circuits. Each 5-HT receptor subtype, 5-HT_1A_R, 5-HT_2A_R, and 5-HT_3A_R, is expressed in a substantial fraction of L5 interneurons, with substantial overlap between subtypes but also cell-type-specific patterns (Santana, Bortolozzi et al. 2004, Puig and Gulledge 2011). As a result, serotonergic modulation in INs can be either excitatory via 5-HT_2A_Rs and 5-HT_3A_Rs or inhibitory via 5-HT_1A_Rs.

Our results demonstrate that 5-HT modulation of L5 INs is highly cell-type specific. Non–fast-spiking (nFS) INs were unresponsive to 5-HT application, consistent with previous reports indicating low or absent expression of serotonergic receptors in this interneuron population (Puig, Santana et al. 2004, Weber and Andrade 2010, Mengod, Palacios et al. 2015). In contrast, both regular fast-spiking (rFS) and stuttering fast-spiking (sFS) INs exhibited a robust 5-HT induced depolarisation. This depolarisation was not blocked by the 5-HT_2A_R antagonist ketanserin, suggesting that 5-HT_2A_Rs are not the primary mediators. Instead, application of the selective 5-HT_3A_R agonist mCPBG evoked a depolarisation, indicating that functional ionotropic 5-HT_3A_R channels are present in this interneuron type. 5-HT_3A_ receptor immunoreactivity has traditionally been associated with nFS interneuron classes such as VIP⁺ and neurogliaform cells (Lee, Hjerling-Leffler et al. 2010, Rudy, Fishell et al. 2011). However, recent transcriptomic and electrophysiological evidence indicates that the population of neurons expressing 5-HT_3A_Rs is more heterogeneous than previously thought (Fuzik, Zeisel et al. 2016, Gouwens, Sorensen et al. 2020). In line with this, our results suggest the presence of functional 5-HT_3A_R-mediated currents in a subset of FS interneurons, revealing an under-appreciated diversity in serotonergic signaling within inhibitory microcircuits. Only a minority of sFS INs showed mild hyperpolarisation upon 5-HT application, which could potentially reflect a concurrent or even dominant expression of inhibitory 5-HT_1A_Rs in these cells. However, this variability was not associated with any clear morphological differences among sFS subtypes. In summary, these findings highlight the functional heterogeneity of 5-HT modulation across electrophysiologically distinct L5 IN subtypes. They underscore the significance of the expression profile and intracellular localisation of 5-HT and other neuromodulator receptors in determining their net effect.

### Presynaptic Modification of Excitatory and Inhibitory Transmission by Serotonin

Apart from its influence on postsynaptic excitability, 5-HT also exerts a powerful presynaptic control over synaptic transmission within local cortical microcircuits. In excitatory-to-excitatory (E–E) connections in layer 5 of the rat mPFC, we observed that 5-HT significantly reduced the amplitude of evoked EPSPs in neuron pairs where the presynaptic neuron was either an AS_H_ or RS pyramidal neuron. In addition, the paired-pulse ratio, coefficient of variation, and failure rate - surrogate measures of synaptic release probability - were also reduced. This suppression was abolished by application of the selective 5-HT_1B_ receptor antagonist SB, indicating that the effect was mediated by 5-HT_1B_Rs. This is in accordance with previous reports of serotonergic suppression of synaptic transmission via presynaptic 5-HT_1B_Rs in visual cortical L5 PNs (Murakoshi, Song et al. 2001), somatosensory cortical L4 neurons (Laurent, Goaillard et al. 2002), and in mouse mPFC L5 PNs (Kjaerby, Athilingam et al. 2016, Berthoux, Barre et al. 2019). Although the literature on E–I modulation is more limited, our findings are consistent with earlier results from the CA1 region of the hippocampus showing that 5-HT_1B_Rs can mediate presynaptic inhibition at both E–E and E–I pathways (Mlinar, Falsini et al. 2003). Together, these data support the idea that 5-HT_1B_Rs suppress synaptic transmission at PN terminals across multiple postsynaptic targets.

In order to investigate by which mechanism 5-HT modulates inhibitory connections at the presynaptic terminal, we recorded cell pairs consisting of presynaptic INs and postsynaptic PNs. In pairs formed by presynaptic sFS and rFS INs, respectively, application of the selective 5-HT_3A_R agonist mCPBG resulted in a significant increase in IPSP amplitude, along with decreased PPR and CV, indicating an increase of GABA release probability. Previous studies reported that 5-HT_3A_Rs, as calcium-permeable ligand-gated cation channels, can enhance presynaptic GABA release (Yang, Mathie et al. 1992, Brown, Hope et al. 1998), and are expressed on a subset of cortical interneurons (Naka and Adesnik 2016, Posluszny 2019). The somatodendritic expression and excitatory function of 5-HT_3A_Rs have been demonstrated previously (Lee, Hjerling-Leffler et al. 2010), while additional studies suggest the localisation of 5-HT_3A_Rs to axon terminal of hippocampal INs (Turner, Mokler et al. 2004, Huang, Yoon et al. 2016).

Our findings indicate that 5-HT_3A_Rs also operate presynaptically in a subset of FS interneurons, a feature that has not been described previously. This observation contrasts with the prevailing view that 5-HT_3A_R expression in the cortex is largely restricted to non-fast-spiking, typically VIP-expressing interneurons (Lee, Hjerling-Leffler et al. 2010, Rudy, Fishell et al. 2011, Naka and Adesnik 2016). These neurons are mainly distributed in layers 2/3, with their density being substantially reduced in the deeper cortical layers. However, most previous studies have not directly examined the functional modulation of synaptic transmission by 5-HT_3A_ receptors. It is therefore possible that presynaptic 5-HT₃A receptor signalling in FS interneurons has been overlooked due to methodological constraints. It should be noted that presynaptic nFS–PN pairs were not recorded in our dataset. This is most likely due to the fact these connections are typically weak and sparse, which renders them less likely to be detected under paired-recording conditions (Xiang, Huguenard et al. 2002, Pfeffer, Xue et al. 2013). Overall, data on presynaptic 5-HT₃A receptor function in layer 5 remain limited, underscoring the need for further experimental investigation.

Taken together, our findings reveal an opposing modulatory role of 5-HT at excitatory and inhibitory synapses: it suppresses glutamatergic transmission via presynaptic 5-HT_1B_Rs, while enhancing GABAergic output from FS interneurons through 5-HT_3A_Rs. This newly identified bidirectional control likely contributes to a net reduction in cortical excitability and reflects a previously unrecognized circuit-level mechanism by which 5-HT shapes local network dynamics. Such opposing actions at distinct synapse types may be critical for mediating the neuromodulatory roles of 5-HT in attention, affect, and cognitive flexibility. Importantly, these effects are mediated by receptor subtypes with distinct cellular and sub-cellular distributions and signalling properties, underscoring the target-specificity of serotonergic control.

## Acknowledgement

We would like to thank Werner Hucko for excellent technical assistance. We thank Dr. Karlijn van Aerde for custom-written macros in Igor Pro software. We are grateful for funding support from the European Union’s Horizon 2020 Framework Programme for Research and Innovation under the Framework Partnership Agreement No. 650003 (HBP FPA) to D.F.

## Data Availability

The datasets generated and analyzed during the current study are available from the corresponding author on reasonable request.

