## Supplemental Table 1 for "Serotonin Engages Divergent 5-HT Receptor Pathways for Cell Type-Resolved Modulation of Prefrontal Layer 5 Microcircuits"

| PSP properties | Amplitude (mV) | | PPR | | Rise time (ms) | | Decay time (ms) | | Latency (ms) | | CV | | Failure rate (%) | |
| --- | --- | --- | --- | --- | --- | --- | --- | --- | --- | --- | --- | --- | --- | --- |
| AS-RS  (n = 10) | 1.1 ± 0.6 | | 1.0 ± 0.2 | | 1.6 ± 1.4 | | 70.2 ± 2.4 | | - | | 0.4 ± 0.2 | | 11.8 ± 10.7 | |
| RS-RS  (n = 8) | 0.8 ± 0.6 | | 0.9 ± 0.3 | | 1.8 ± 1.1 | | 71.8 ± 2.5 | | - | | 0.5 ± 0.2 | | 16.3 ± 15.4 | |
| P-value | ns | 0.35 | ns | 0.20 | ns | 0.78 | ns | 0.11 | - | | ns | 0.41 | ns | 0.46 |
| AS-bFS  (n = 6) | 3.5 ± 3.6 | | 0.8 ± 0.2 | | 0.7 ± 0.4 | | 67.3 ± 3.7 | | 1.0 ± 0.4 | | 0.7 ± 0.5 | | 10 ± 20 | |
| RS-rFS  (n = 8) | 1.9 ± 1.5 | | 0.8 ± 0.04 | | 0.8 ± 0.3 | | 65.7 ± 7.1 | | 1.4 ± 1.4 | | 0.4 ± 0.2 | | 20 ± 20 | |
| P-value | ns | 0.29 | ns | 0.33 | ns | 0.67 | ns | 0.57 | ns | 0.53 | ns | 0.84 | ns | 0.88 |
| bFS-AS  (n = 6) | 0.7 ± 0.5 | | 0.8 ± 0.2 | | 1.4 ± 0.2 | | 64.1 ± 8.5 | | 0.9 ± 0.2 | | 0.5 ± 0.1 | | 10.9 ± 13.7 | |

**Table S1: Unitary EPSP characteristics of different subtypes of L5 excitatory and inhibitory connections.** The postsynaptic properties are shown as average ± SD. For pairs recorded in the ‘loose-seal’ configuration, the latency was not calculated.

| PSP properties | E-E (n = 15) | | | | | | | | | E-I (n = 12) | | | | I-E (n = 7) | | | |
| --- | --- | --- | --- | --- | --- | --- | --- | --- | --- | --- | --- | --- | --- | --- | --- | --- | --- |
|  | CTL | 5-HT | SB | CTL vs 5-HT | | 5-HT vs SB | | CTLvs SB | | CTL | CGS | CTL vs CGS | | CTL | mCPG | CTL vs mCPG | |
| Amplitude (mV) | 1.0 ± 0.6 | 0.6 ± 0.3 | 0.8 ± 0.4 | *** | 6,1E-05 | *** | 6,1E-05 | ns | 0,07 | 2.2 ± 2.6 | 1.5 ± 2.1 | ** | 0,0005 | 0. 5 ± 0.5 | 1.0 ± 0.7 | * | 0,03 |
| PPR | 0.9 ± 0.2 | 1.1 ± 0.3 | 0.9 ± 0.3 | *** | 6,1E-05 | *** | 6,1E-05 | ns | 0,82 | 0.7 ± 0.2 | 1.5 ± 0.7 | ** | 0,0005 | 0.9 ± 0.1 | 0.7 ± 0.1 | * | 0,03 |
| Rise time (ms) | 1.9 ± 1.3 | 1.6 ± 0.9 | 1.6 ± 0.7 | ns | 0,16 | ns | 0,39 | ns | 0,33 | 0.7 ± 0.3 | 0.7 ± 0.3 | ns | 0,40 | 1.8 ± 0.6 | 2.3 ± 0.9 | * | 0,03 |
| Decay time (ms) | 70.9± 2.6 | 71.2 ± 3.0 | 71. 3 ± 3.2 | ns | 0,23 | ns | 0,50 | ns | 0,97 | 66.7 ± 4.6 | 68.1 ± 5.0 | ns | 0,11 | 65.6 ± 9.2 | 60.2 ± 8.8 | * | 0,03 |
| Latency (ms) | - | - | - |  | - |  | - |  | - | 1.4 ± 1.1 | 1.7 ± 1.1 | * | 0,005 | 0.9 ± 0.2 | 0.7 ± 0.3 | ns | 0,94 |
| CV | 0.4 ± 0.2 | 0.7 ± 0.3 | 0.5 ± 0.3 | *** | 6,1E-05 | *** | 0,0001 | ns | 0,49 | 0.7 ± 0.4 | 0.9 ± 0.4 | ** | 0,0005 | 0.6 ± 0.1 | 0.4 ± 0.2 | * | 0,03 |
| Failure rate (%) | 11.2 ± 10.5 | 31.8 ± 22.3 | 16.7 ± 20.0 | ** | 0,001 | ns | 0,09 | ns | 0,18 | 22.0 ± 20.5 | 34.3 ± 24.1 | ** | 0,0005 | 18.3 ± 19.0 | 7.6 ± 9.6 | ns | 0,06 |

**Table S2: PSP characteristics of different subtypes of L5 excitatory and inhibitory connections.** The postsynaptic properties are shown as average ± SD. For pairs recorded in the ‘loose-seal’ configuration, the latency was not calculated. SB5: 5-HT_1B_R antagonist (5 µM); CGS: 5-HT_1B_R agonist (5 µM); mCPG: 5-HT_3A_R agonist (30 µM).
